## Supplemental Figures and Text for "Dock-and-lock binding of SxIP ligands is required for stable and selective EB1 interactions"

**Supplementary information**

**Supplementary Figures**

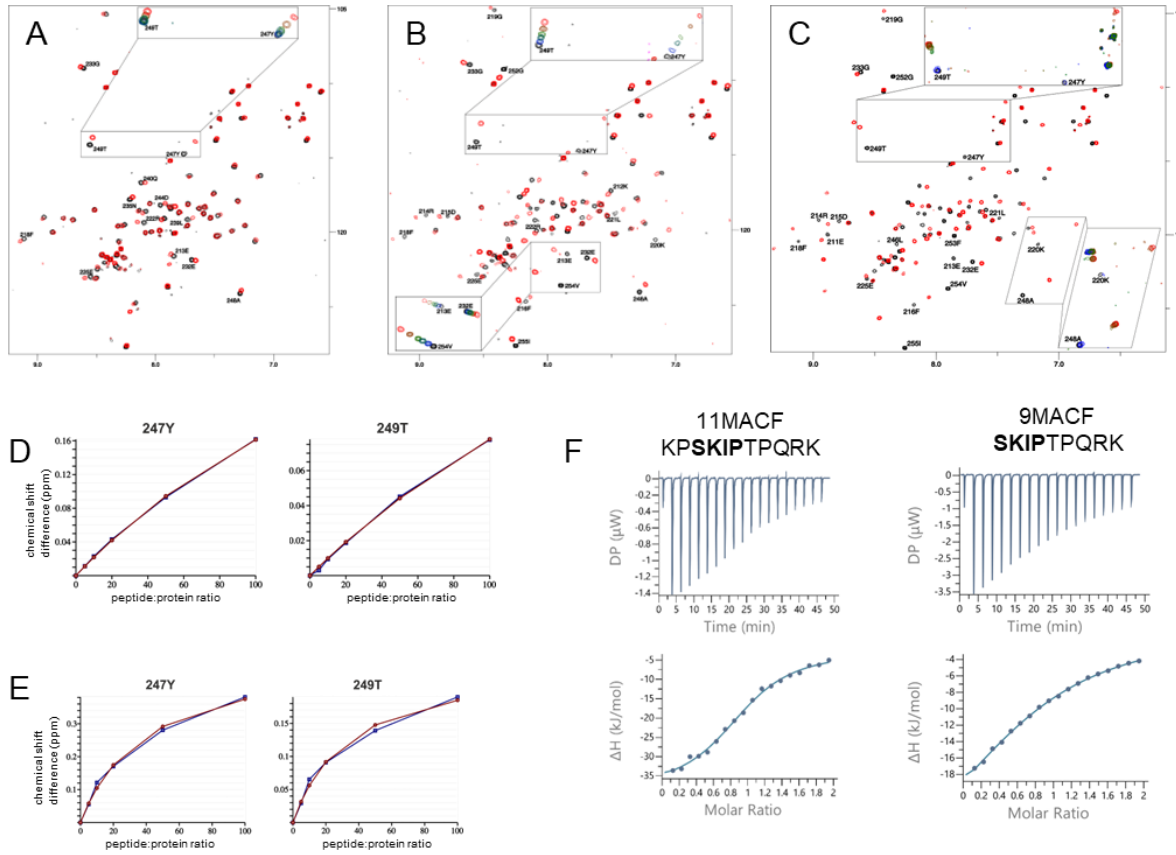


**Figure S1**. Post-SxIP residues increase the binding affinity. (A, B and C) Superposition of $^{1}$H,$^{15}$N-HSQC spectra of $^{15}$N-labelled EBH domain of EB1 (50 µM) recorded at 600 MHz for the free form (black) and in the presence of the peptide (black), (A) 4MACF (5000 µM), (B) 6MACF (5000 µM), and (C) 11MACF (400 µM). The inserts show changes in the spectra on the increase of the peptide concentration, (A) 250, 500, 1000, 2500 and 5000 µM of 4MACF, (B) 250, 500, 1000, 2500 and 5000 µM of 6MACF, and (C) 12.5, 25, 75, 82.5, 100, 125, 150, 200 and 400 µM of 11MACF. (D and E) Chemical shift changes (blue) of the residues with the largest induced shifts fitted into the single site binding model (red) to evaluate the dissociation constant $K_D$ for (D) 4MACF and (E) 6MACF. (F) ITC isotherm for the interaction of EBH with 11MACF (left) and 9MACF (right). The solid line shows the fitting of the data into a single-site binding model.

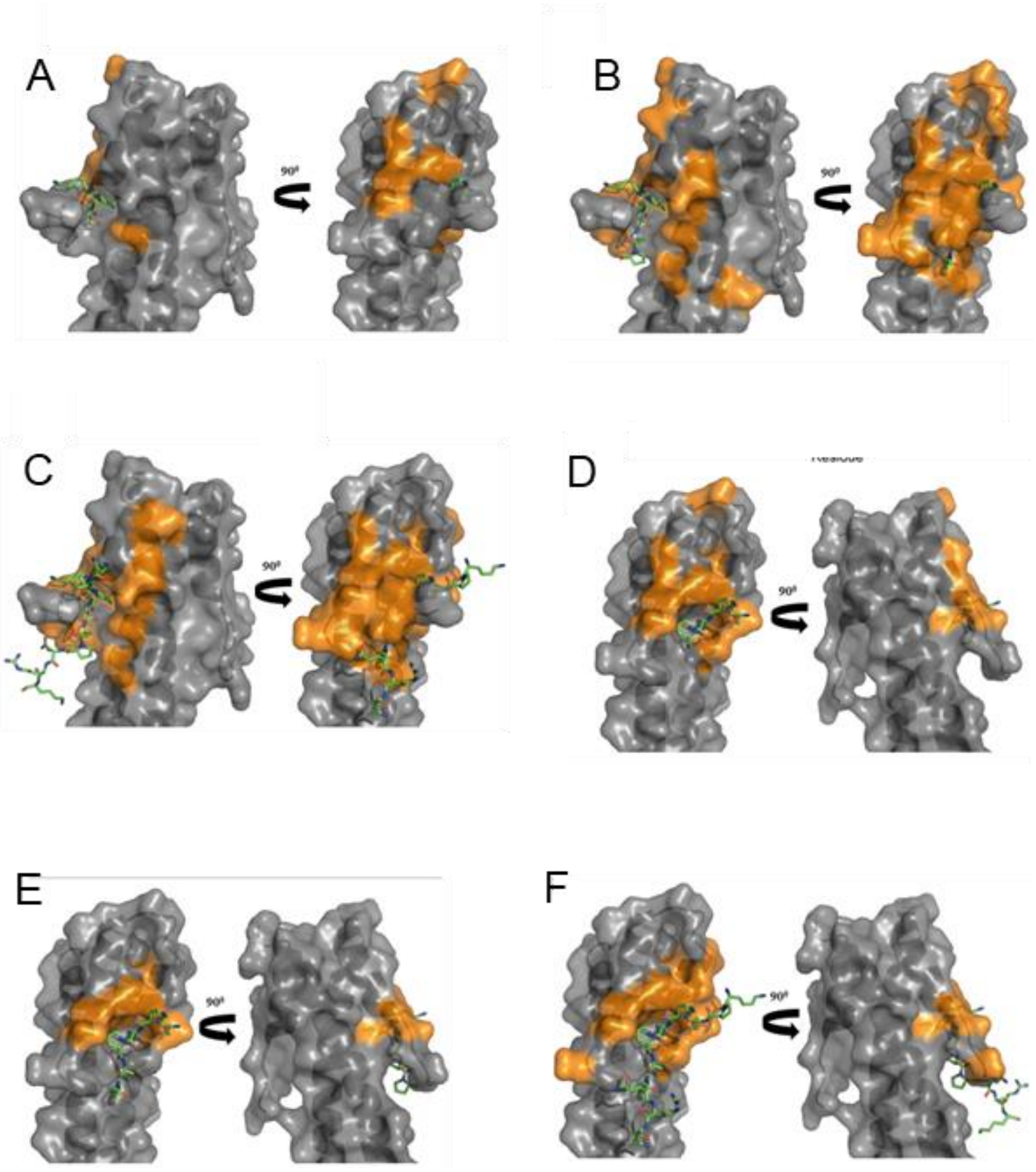


**Figure S2**. Mapping of the largest chemical shift changes induced by the peptide binding on the surface of EB1 EBH domain (A-C) and EBH EBH-$\Delta$C (D-F) in the complex with the MACF peptide (green, shown in a stick representation). The crystal structure was used to represent the complex (PDB ID 3GJO). The EBH C-terminus has been removed for EBH$\Delta$C. (A-C) Changes induced in the $^{1}$H,$^{15}$N-HSQC signals of EBH by (A) 4MACF, (B) 6MACF, and (C) 11MACF peptides. (D-F) Changes induced in the $^{1}$H,$^{15}$N-HSQC signals of EBH$\Delta$C by (D) 4MACF, (E) 6MACF, and (F) 11MACF peptides.

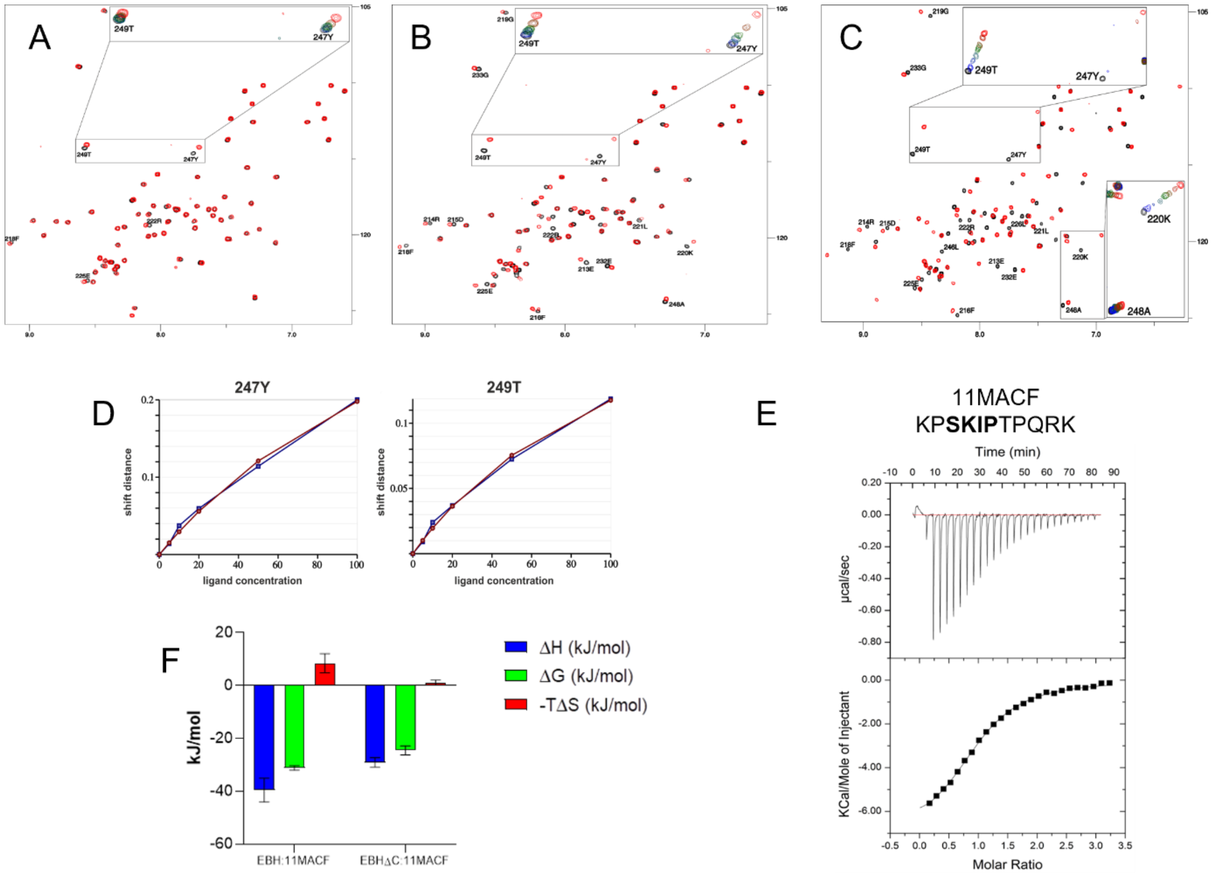


**Figure S3**. Deletion of the EBH C-terminus reduces the binding affinity. (A, B and C) Superposition of $^{1}H$,$^{15}N$-HSQC spectra of $^{15}N$-labelled EBH-ΔC domain of EB1 (50 µM) recorded at 600 MHz for the free form (black) and in the presence of the peptide (black), (A) 4MACF (5000 µM), (B) 6MACF (5000 µM), and (C) 11MACF (400 µM). The inserts show changes in the spectra on the increase of the peptide concentration, (A) 250, 500, 1000, 2500 and 5000 µM of 4MACF, (B) 250, 500, 1000, 2500 and 5000 µM of 6MACF, and (C) 12.5, 25, 75, 82.5, 100, 125, 150, 200 and 400 µM of 11MACF. (D) Chemical shift changes (blue) of the residues with the largest induced shifts fitted into a single-site binding model (red) to evaluate the dissociation constant $K_D$ for 6MACF. (E) ITC isotherm for the interaction of EBH with 11MACF. The solid line shows the fitting of the data into a single-site binding model. (F) Thermodynamic parameters (ΔG, blue; ΔH, red; -TΔS, green) evaluated from the ITC data for the 11MACF binding to the EBH (left) and EBHΔC (right).

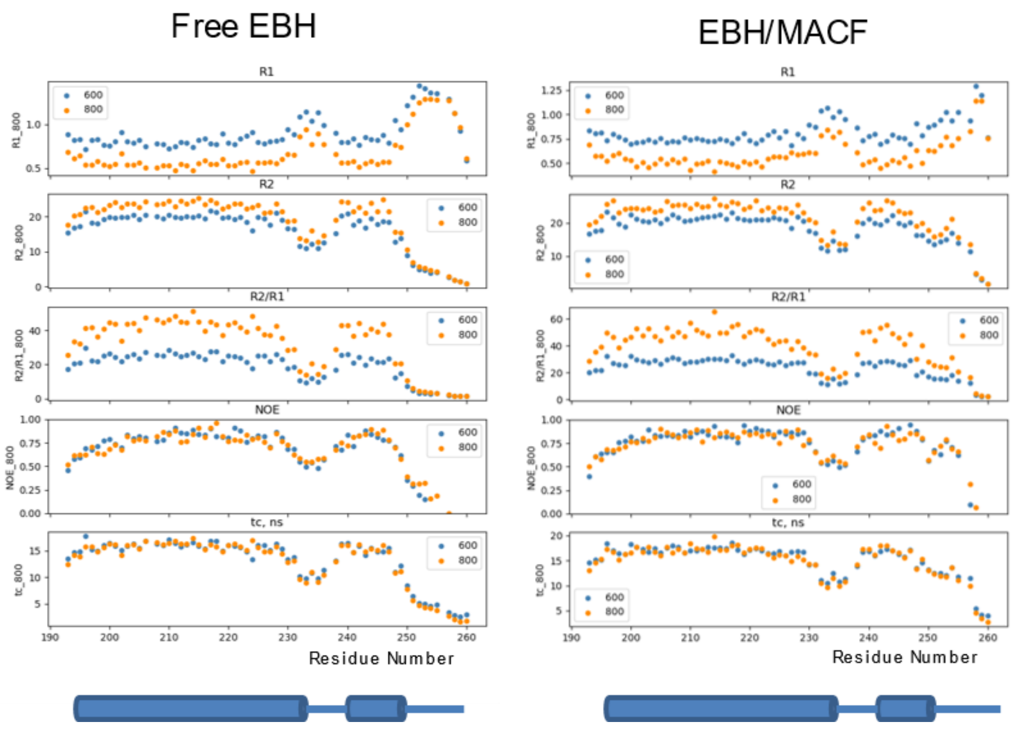


**Figure S4**. Relaxation parameters measure at 600 (blue) and 800 (orange) MHz for free EBH (left) and EBH in the complex with 11MACF (right). From top to bottom: relaxation rates R1, R2, ratio R2/R1, $^{1}H$-$^{15}N$ NOE, and isotropic correlation time $\tau_c^{iso}$ directly calculated for each residue using isotropic model. The secondary structure of the EBH domain is shown schematically at the bottom.

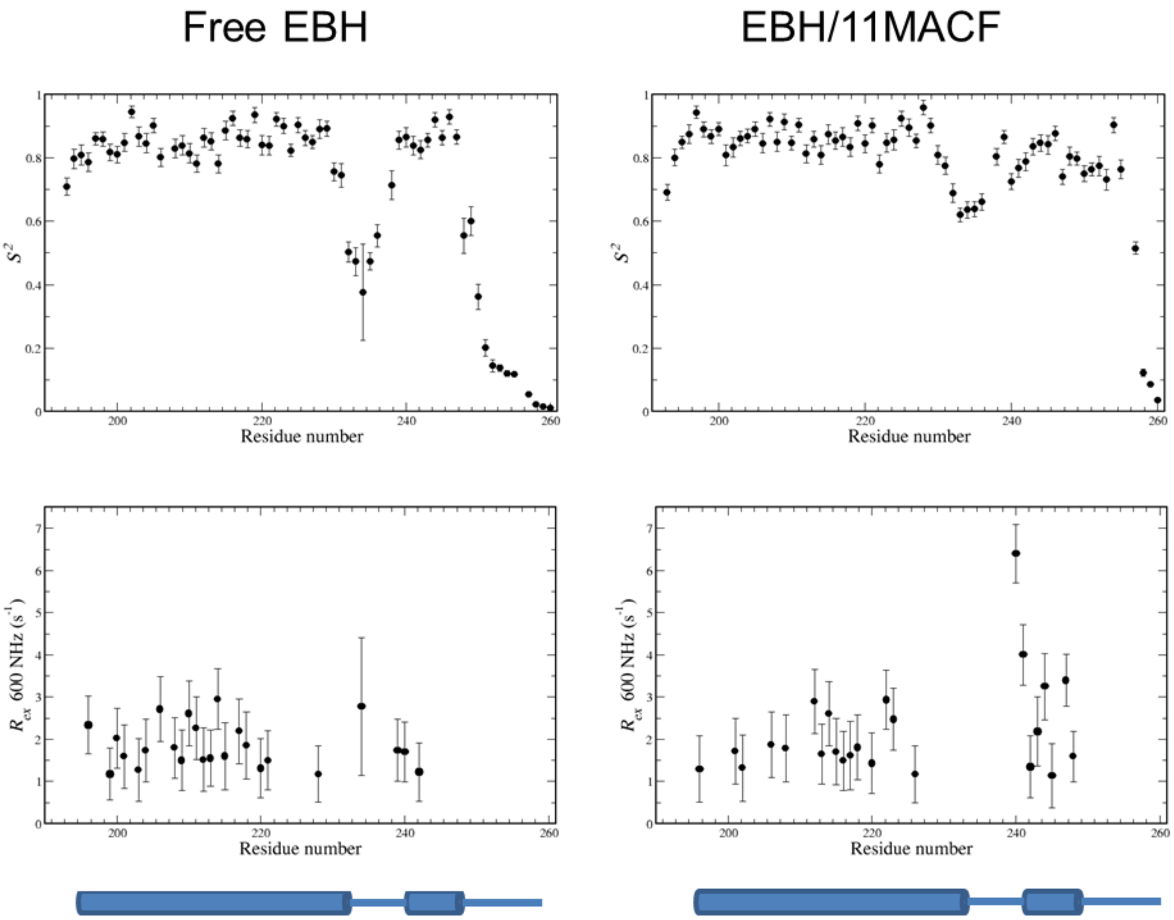


**Figure S5**. Order parameters $S^2$ and exchange rates $R_{ex}$ calculated from the relaxation data for free EBH (left) and EBH in complex with the 11MACF peptide (right).

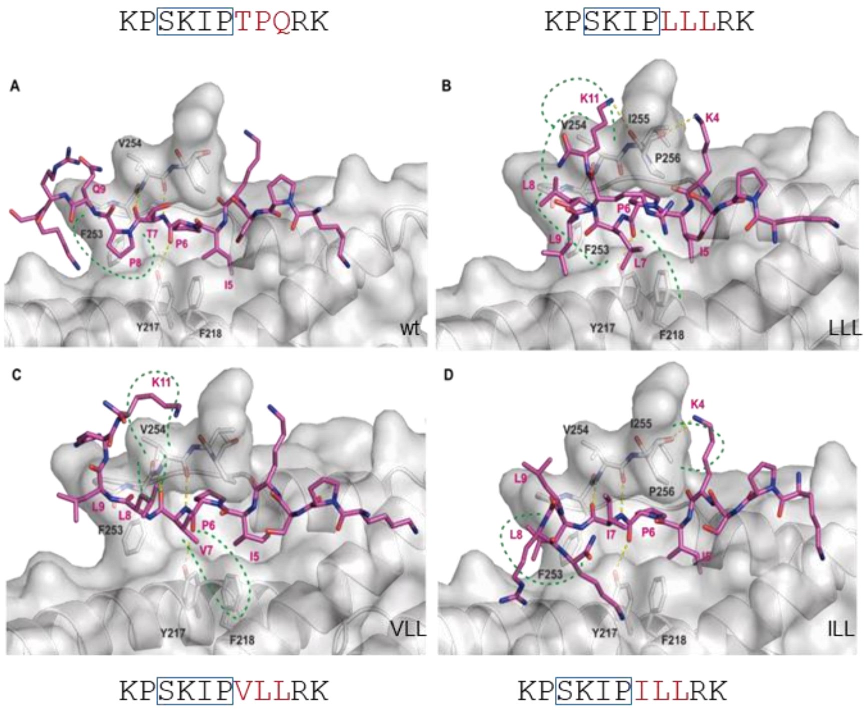


**Figure S6**. Best scored docking poses obtained for (A) – 11MACF-WT, (B) – 11MACF- LLL, (C) - 11MACF-VLL and (D) – 11MACF-ILL. The EBH structure is shown as a semi-transparent grey surface and cartoon. The peptide is shown in stick representation. Carbon atoms are coloured in magenta, oxygen is shown in red and nitrogen in blue. Hydrogen bonds are shown as yellow dashed lines and hydrophobic interactions regions are highlighted by green dashed lines. The SxIP region is boxed.

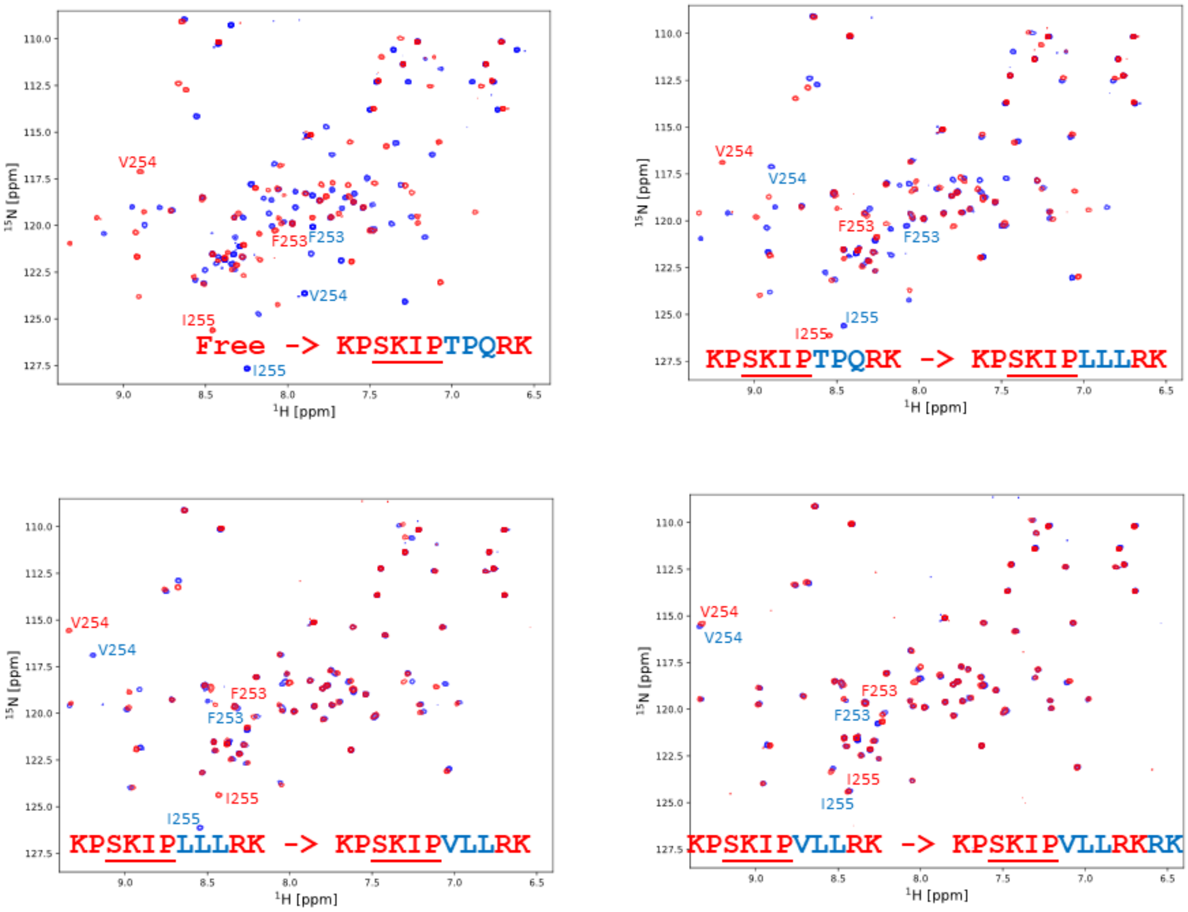


**Figure S7**. Progressive changes in the HSQC spectra in the complexes with the mutant peptides. Superposition of (A) the $^{1}$H,$^{15}$N-HSQC spectra of free EBH (blue) and EB1/11MACF complex (red), (B) EB1/11MACF (blue) and EB1/11MACF-LLL (red) complex, (C) EB1/11MACF-LLL (blue) and EB1/11MACF-VLL (red) complex, and (D) EB1/11MACF-VLL (blue) and EB1/11MACF-VLLRK (red) complex. The SxIP region is underlined and the modified post-SxIP region is highlighted in blue. Signals corresponding to the $^{253}$FVI$^{255}$ region are labelled in the spectra, with colours corresponding to the spectrum.

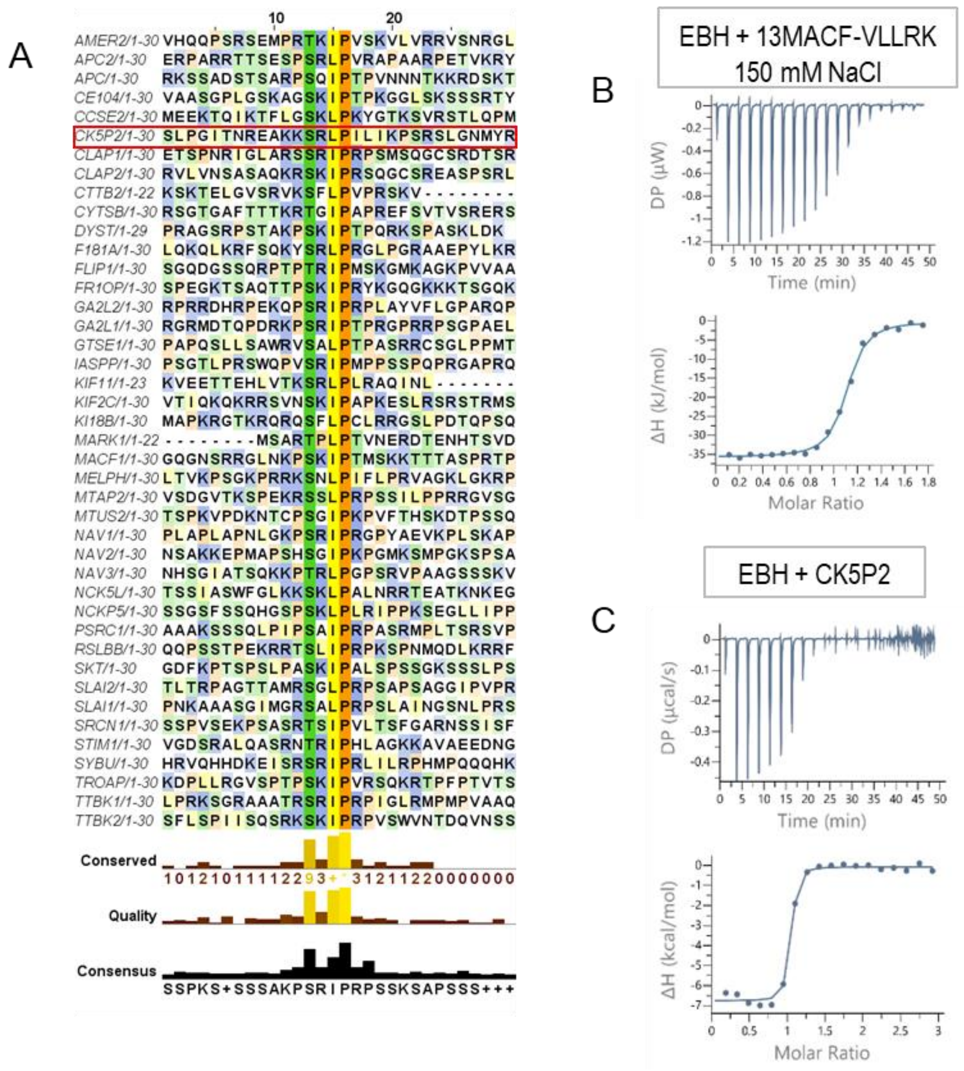


**Figure S8**. (A) Sequence alignment of known SxIP proteins showing large variation of non-SxIP residues. Sequence conservation and consensus sequence are presented at the bottom. Residues are colour coded according to the properties of the side-chains. The figure was created with JalView. (B, C) ITC isotherm for the interaction of EBH with 13MACF (B) and CK5P2 (C). The solid line shows the fitting of the data into a single-site binding model.

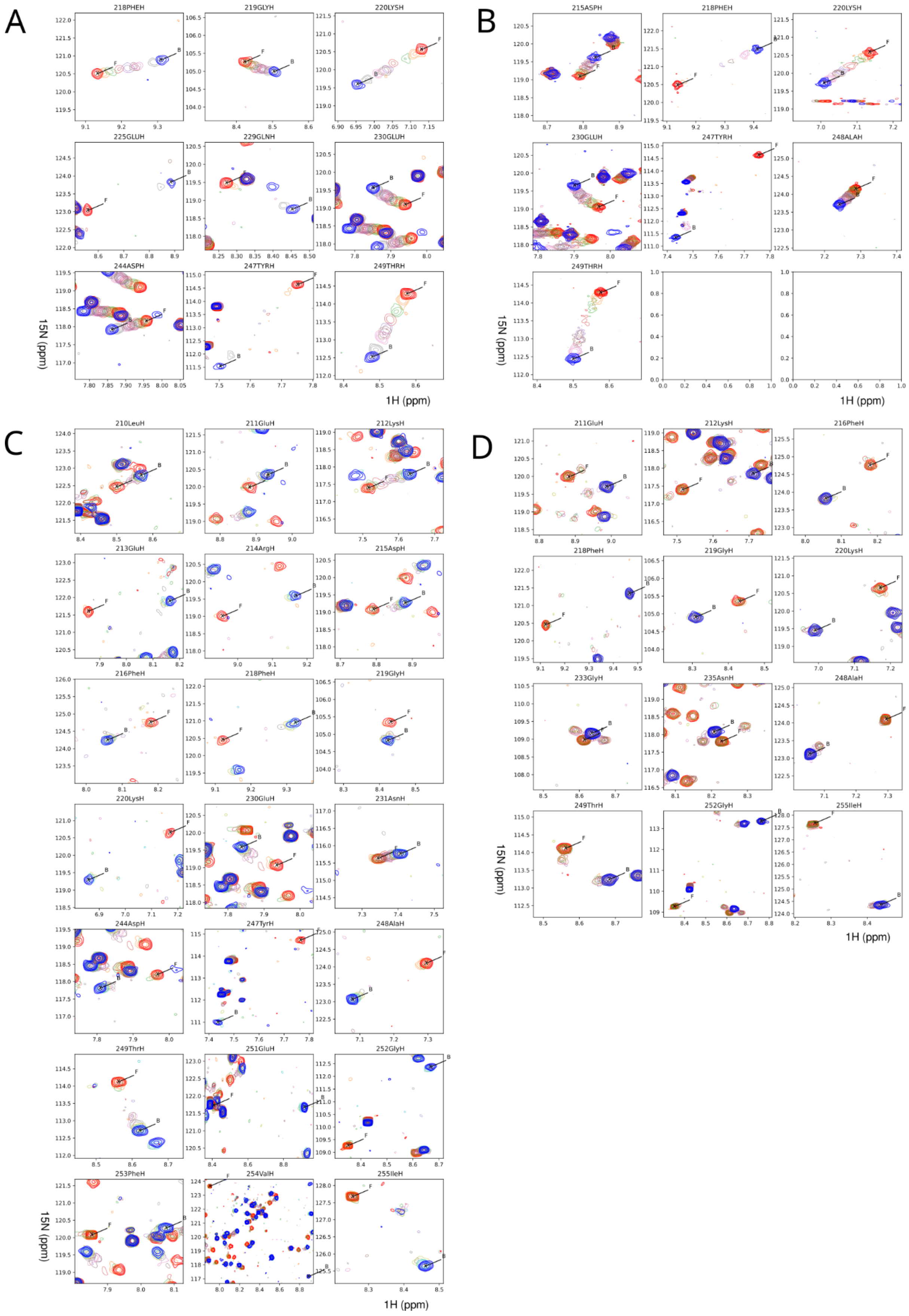
A
218PHEH
219GLYH
220LYSH
225GLUH
229GLNH
230GLUH
244ASPH
247TYRH
249THRH
15N (ppm)
1H (ppm)
B
215ASPH
218PHEH
220LYSH
230GLUH
247TYRH
248ALAH
249THRH
15N (ppm)
1H (ppm)
C
210LeuH
211GluH
212LysH
213GluH
214ArgH
215AspH
216PheH
218PheH
219GlyH
220LysH
230GluH
231AsnH
244AspH
247TyrH
248AlaH
249ThrH
251GluH
252GlyH
253PheH
254ValH
255IleH
15N (ppm)
1H (ppm)
D
211GluH
212LysH
216PheH
218PheH
219GlyH
220LysH
233GlyH
235AsnH
248AlaH
249ThrH
252GlyH
255IleH
15N (ppm)
1H (ppm)

**Figure S9**. Progressive changes in the $^1H,^{15}N$-HSQC spectra on peptide addition observed for different EB1 interactions that were used to evaluate the exchange rates with TITAN software for the corresponding EBH complexes. Superposition of the spectra for the titration of (A) EBH-ΔC with 11MACF, (B) EBH-ΔC with 11MACF-VLL, (C) EBH with 11MACF, and (D) EBH with 11MACF-VLL. Signals of the free EBH are shown red, final titration point in blue, and the intermediate titration points are shown in the pale colours. Notice additional signals that are observed for EBH/11MACF-VLL titration at the intermediate concentrations corresponding to the non-symmetrical form where the EBH dimer binds a single peptide. These signals can only be observed when the exchange between different forms is very slow.

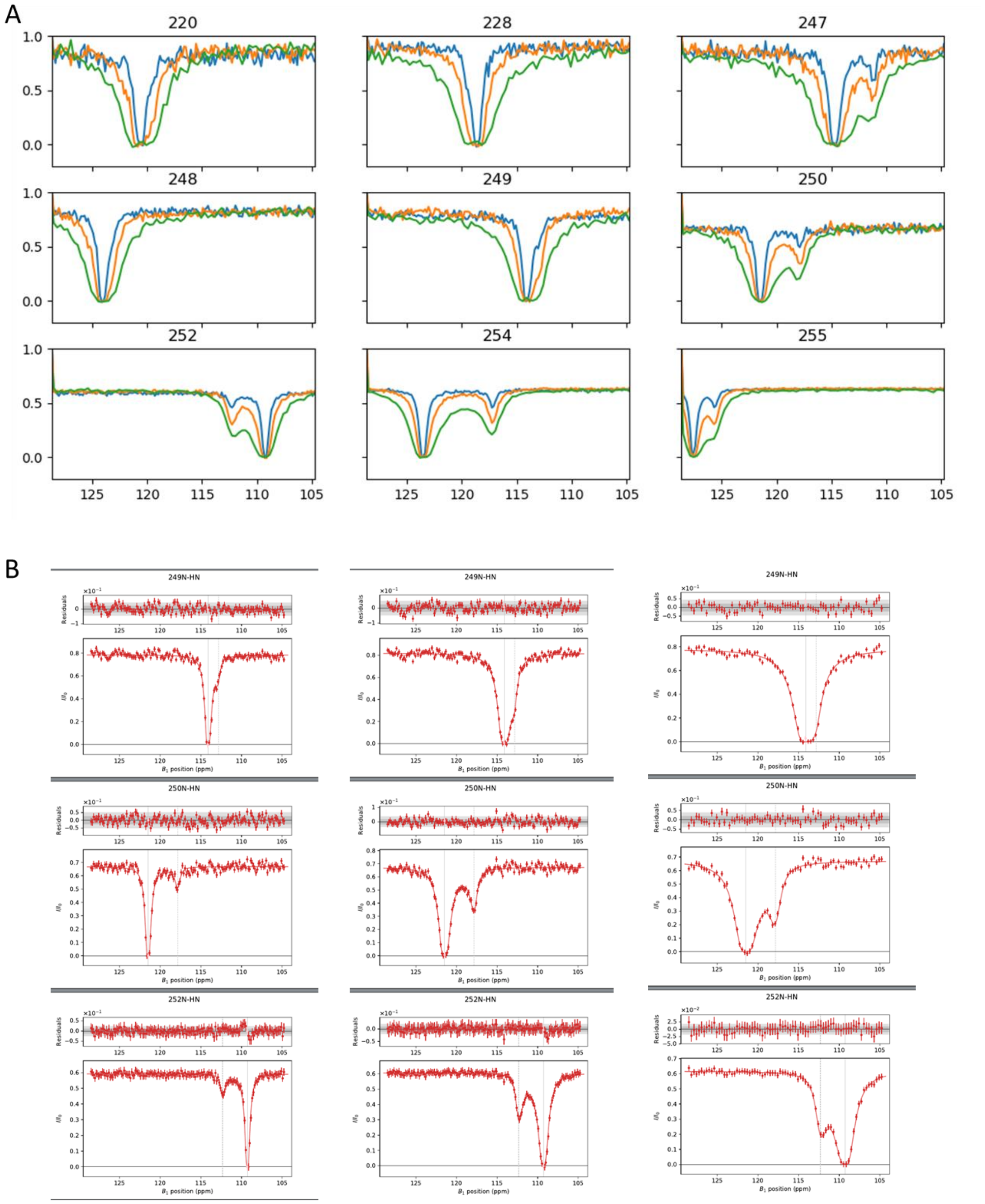


Figure S9

**Figure S10**. (A) Summary of the CEST profiles observed for EBH/11MACF interaction at the irradiation field strength of 12.5 (blue), 25 (orange) and 50 Hz (green). (B) Illustration of the CEST profile fitting (solid line) into the global exchange model with the dissociation rate 120 $s^{-1}$ calculate with Chemix software.

**Supplementary text**

**TITAN models for fitting EBH titration data.**

In general, the dimeric structure of EBH and the location of the binding site at the interface between the monomers requires the use of the dimer binding model with three different states: free EBH, EBH bound to one peptide and the full complex with two peptides bound. In both the free and the final complex, the chemical shifts of both subunits are identical due to the symmetry of the EBH dimer. However, binding to only one site may have an indirect effect on the chemical shifts of the other site and the appearance of two additional signals at the intermediate points of the titration; one corresponding to the empty, and one to the filled site that are different to the shifts of the free and fully bound states. The indirect effects on the chemical shits are usually small, leading to the cross peaks of the empty site being close to the free state, and the occupied site to the peaks of the fully bound state. These additional cross-peaks can only be observed if the exchange rate is sufficiently slow. In the case of the fast exchange the presence of the intermediate state can be detected as a complex, non-linear pattern of the chemical shifts (Waudby, Ramos et al. 2016). In such cases a 4-state model is required for the fitting. However, if the indirect effect on the chemical shifts is small and the exchange is fast, the binding to each site can be treated as independent and the changes in the spectra can be fitted with a two-state binding model.

We observed different exchange regime for the interactions of EBH and EBH-ΔC with different MACF peptides, in agreement with the dissociation constants (Table 1). The interaction of the 11MACF with EBH-ΔC (ITC $K_D$ 41.5 ± 8.8 uM) showed gradual chemical shift changes with a significant peak broadening for the largest chemical shift differences, corresponding to the fast-intermediate exchange (Figure S8A). The majority of the peak trajectories followed a straight line, in agreement with small indirect binding effect and applicability of the two-site binding model. The TITAN analysis with the single site model (independent binding to the two binding sites of the dimer) gave $K_D = 26.6 \pm 0.051\ \mu M$ that agreed well with the ITC $K_D$, and $k_{off} = 1900 \pm 18\ s^{-1}$, corresponding to the fast exchange. The mutant 11MACF-VLL peptide showed stronger binding by ITC (ITC $K_D$ 18.7 ± 3.1 uM). In agreement with this we observed more pronounced broadening during the NMR titration (Supplementary Figure S9B). TITAN analysis of the data gave $K_D = 25.62 \pm 0.82\ \mu M$ that agreed well with the ITC $K_D$, and $k_{off} = 1541 \pm 79\ s^{-1}$, that was close to the off-rate of the WT peptide. This demonstrated that the affinity increase was caused mainly by the on-rate increase.

For the significantly stronger interaction of the 11MACF with EBH (ITC $K_D$ 3.5 ± 1.0 uM) we observed a combination of intermediate exchange for smaller chemical shift changes and slow exchange for the largest changes in signals of the residues in the binding pocket, particularly in the C-terminal region that becomes immobilised in the complex (Figure S8B). In a number of cases the peak trajectories were not linear, indicating an indirect effect on chemical shifts in one site from the binding of the peptide to the other site. At the lower temperature of 15$^{o}$C additional peaks were observed for the intermediate binding state. These signals correspond to the non-symmetrical EBH complex where one binding site is occupied, and the other binding site is empty. Although the two binding sites are located at the opposite sides of the EBH dimer and do not overlap, the peptide binding to one site causes small changes in the EBH structure propagated to the un-occupied site. This, in turn, leads to the small chemical shift changes in the empty site. In a similar way, the binding of the second peptide causes small structural changes in the occupied site, and, thus, additional chemical shift changes. As the result, in the non-symmetrical complex with one peptide bound, the signals of free site are different from the signals of the free EBH, and the signals of the bound site are different from the signals of the symmetrical complex with the two peptides bound. This leads to the appearance of additional signals in the intermediate bound state when the exchange is slow, or non-linear signal behaviour under sufficiently fast exchange condition. The additional signals confirm the binding to the EBH dimer. The small value of the chemical shift changes induced by the indirect effect show that the structural changes are very small.

In the case of non-linear chemical shift changes or appearance of new signals, the single-side binding model cannot reproduce the spectral variations and the dimer 4-state binding model with additional parameters has to be used for the fitting. The dimer binding model in TITAN includes cooperativity factors $\alpha = \log(K_{D,2}/K_D)$ and $\beta = \log(k_{off,2}/k_{off})$, where $K_{D,2}$ and $k_{off,2}$ are the dissociation constant and off-rate of the second binding, respectively. The fitting of the titration with all parameters showed close to zero values for both cooperativity parameters. We, therefore, fixed their values at zero during the fit to simplify the model. The resulting values of $K_D = 4.9 \pm 0.1\ \mu M$ and $k_{off} = 130.2 \pm 2.1\ s^{-1}$ that were close to the values obtained with the cooperativity parameters ($K_D = 4.8 \pm 0.1\ \mu M$ and $k_{off} = 130 \pm 0.7\ s^{-1}$).

For the stronger interaction between 11MACF-VLL and EBH (ITC Kd 80 nM) many signals showed slow exchange and clear presence of signals from the intermediate binding state (Figure S8C), which required the use of the dimer binding model. In most cases one of the intermediate state signals was located close to the free, and the other close to the fully bound-state signal, in agreement with a much smaller indirect effect. However, for the residues in the loop between the short and the long helix Gly230, Asn231, Gly233 and Asn235 signals of the intermediate state were significantly more displaced from the signals of the free and fully bond-forms (Figure S8C). These residues are located far from the binding site and do not make direct contact with the peptide. The effect on the chemical shifts of these residues indicates some structural rearrangement of the relative positions of the helices induced by the peptide binding to each of the sites.

The fitting of the 11MACF-VLL titration with all for parameters resulted in $K_D = 66 \pm 2\ nM$, $k_{off} = 15 \pm 0.2\ s^{-1}$, $\alpha = 0.5$ and $\beta = 0$, corresponding to a weak negative cooperativity with a ~3-fold increase of the $K_D$ for the second binding without a change in the dissociations rate. The dissociation constant for the first binding event agreed very well with the ITC dissociation constant of $80 \pm 2\ nM$, supporting the NMR analysis. The negative cooperativity may reflect the small structural rearrangement detected from the chemical shift changes. When the cooperativity parameters were set to zero during the fit, corresponding to the two-state binding model used for ITC, the $K_D$ value increased to $200 \pm 4\ nM$, while the dissociation rate remained unchanged. The lack of the systematic deviations between the experimental ITC data and the single site analysis model shows that the difference between the first and the second site binding, if exists, is too small to be detected by ITC, in line with the weak cooperativity observed by NMR. We therefore concluded that while NMR gives indication of the weak negative cooperativity, the differences in the binding affinities for the first and second ligand binding events are too small for a reliable interpretation and the single site binding model should be used to avoid overfitting and overinterpretation.

**Exchange approximation for low population of the intermediate state.**

From the $K_D'$ and $k_{off}'$ values for the EBH/11MACF interaction, we can estimate $K_D^L = 0.08 -$

0.07, showing that only ~8% of the bound population is in the intermediate dock state. Since the intensities of the NMR signals corresponding to the intermediate state are proportional to the population of this state, its contribution into the spectra is negligibly small compared to the fully bound state and can be neglected. Under this condition, the measured line-shape changes and CEST saturation transfer are only associated with the exchange between the free and the fully bound forms. These rates can be calculated as the overall fluxes between the states using the steady-state conditions as (Shoup and Szabo 1982):

$$k'_{on} = k^D_{on} \frac{k^L_{on}}{k^D_{off} + k^L_{on}} \tag{1}$$

$$k'_{off} = k^D_{off} \frac{k^L_{off}}{k^D_{off} + k^L_{on}} \tag{2}$$

The effective dissociation constant between these states is then:

$$\boldsymbol{K'_D = K^D_D * K^L_D} \tag{3}$$

which agrees with the Equation 1 when $K^D_D \ll 1$.

Equation 3 can be rearranged:

$$k'_{on} = k^D_{on} \frac{1}{\frac{k^D_{off}}{k^L_{on}} + 1} \tag{4}$$

Since:

$$\frac{1}{\frac{k^D_{off}}{k^L_{on}} + 1} \leq 1 \tag{5}$$

then

$$k'_{on} \leq k^D_{on} \tag{6}$$

From this equation, the additional exchange step caused by the conformation change of the complex should generally lead to a reduction in the on-rate for the full transition to the complex. In the case of a very fast transfer from the intermediate to the fully bound state compared to the ligand dissociation from the transition state ($k^D_{off} \ll k^L_{on}$), the initial dock step becomes rate-limited, with overall on-rate equal to the on-rate of the dock step.

From the equations (1) and (2) of the main text,

$$\frac{\boldsymbol{K'_D}}{\boldsymbol{k'_{off}}} = \frac{\boldsymbol{K_D^D}}{\boldsymbol{k_{off}^D}} \tag{7}$$

and

$$k'_{on} = k_{on}^D \tag{8}$$

So, the equality of the on-rates also holds under fast exchange in the second lock step even if the population of the intermediate state is not negligibly small.

When the population of the intermediate state is comparable to the population of the fully bound state and the exchange in the second lock step is intermediate or slow, the binding cannot be approximated by a two-site model and the intermediate step should be considered explicitly. Under these conditions, NMR signals of the fully titrated protein will have an exchange contribution or additional signals corresponding to intermediate state will be observed. This will allow direct evaluation of the exchange parameters of the second lock step from the NMR analysis.

**Table S1 –** NMR restraints and structure statistics for the NMR structure of the EBH/11MACF complex.

| Total restraints used | |
|---|---|
| NOE restraints* | |
| *All* | 3863 |
| *Protein-ligand* | 298 |
| *Intermonomer* | 924 |
| *Intrapeptide* | 209 |
| *Intraresidue* | 1133 |
| *Sequential* ($\|i-j\| = 1$) | 968 |
| *Medium* ($1 < \|i-j\| \leq 4$) | 1311 |
| *Long range* ($\|i-j\| > 4$) | 177 |
| Dihedral | |
| $\phi$ *angles* | 65 |
| $\varphi$ *angles* | 65 |
| Hydrogen bonds | 90 |
| Structure statistics | |
| Violations | |
| *Distance (> 0.5 Å)* | 41 |
| *Dihedral angle* ($> 5^0$) | 7 |
| Energies (cal/mol) | |
| *Overall* | -2179 (±208) |
| *Bond* | 119 (±9) |
| *Angle* | 479 (±21) |
| *Improper* | 242 (±29) |
| *Dihedral* | 868 (±14) |
| *Van der Waals* | -48 (±28) |
| *Electrostatic* | -5903 (±98) |
| *NOE* | 1906 (±119) |
| Geometry – average Values | |
| *Bond* | $7.40 \times 10^{-3}$ ($\pm 5.7 \times 10^{-4}$) |
| *Angle* | 0.91 ($\pm 9.96 \times 10^{-2}$) |
| *Improper* | 2.50 (±0.30) |
| *Dihedral* | 41.56 (±0.24) |
| *Van der Waals* | 428.93 (±83.98) |
| Average pairwise RMSD (Å)** | |
| *Heavy atoms* | 2.41 (±1.25) |
| *Heavy atoms – helical region* | 1.51 (±0.99) |
| *Backbone* | 2.05 (±1.33) |
| *Backbone – helical region* | 1.08 (±0.93) |
| Ramachandran statistics (%) *** | |
| *Most favoured regions* | 87.0 (99.5) |
| *Additional allowed regions* | 10.8 (0.3) |
| *Generously allowed regions* | 1.0 (0.2) |
| *Disallowed regions* | 1.1 (0) |

*Number in brackets corresponds to the restraints assigned manually

**Helical region corresponds to residues: Glu192-Glu230 and Pro237-Tyr247

***Values within brackets correspond to residues Glu192-Glu230 and Pro237-Tyr247 (helical region)
